# From Archaea to the atmosphere: remotely sensing Arctic methane

**DOI:** 10.1101/2025.02.13.638097

**Authors:** Ruth K. Varner, Dylan R. Cronin, Patrick Crill, Michael Palace, Carmody K. McCalley, Jia Deng, Christina Herrick, McKenzie Kuhn, Suzanne B. Hodgkins, Kellen McArthur, Jessica DelGreco Singer, Benjamin Bolduc, Yueh-Fen Li, The Archaea to Atmosphere (A2A) Project Team, EMERGE Institute Coordinators, Changsheng Li, Gene Tyson, Steve Frolking, Jeffrey P. Chanton, Andreas Persson, Scott R. Saleska, Virginia I. Rich

## Abstract

Global atmospheric methane concentrations are rapidly rising and becoming isotopically more depleted, implying an unresolved microbial contribution. Rising Arctic temperatures are variably altering soil methane cycling, causing consequential uncertainty in the atmospheric methane budget. We demonstrated in an Arctic wetland that below-ground microbiota and methane-cycling features parallelled above-ground plant communities. To upscale emissions, we applied machine learning to remote sensing data to identify habitats, which were assigned average emissions. To upscale dynamically, we incorporated climate data, remotely-sensed water table variation, and habitat classes into a temporally-resolved biogeochemical model, to predict methane flux and isotope dynamics. This accurately estimated more depleted 13C-methane than previously used for Arctic habitats in global source partitioning. Remote-sensing of these rapidly changing inaccessible landscapes can thus help constrain the role of the Arctic in ongoing changes in global methane emissions.

## Main text

Accurate estimates of climate-warming gas emissions are essential to understand past climate variability, evaluate ongoing climate change mitigation strategies, and predict future climate trajectories. However, large discrepancies persist between bottom-up (upscaled empirical measurements and process-based models) and top-down (derived from atmospheric inverse modeling) emission estimates, and each has significant uncertainties (Saunois et al. 2020; Ramage et al. 2024). One approach to the former is landcover-class scaling: developing landcover products (maps using geospatial data and remote-sensing data from satellites; e.g. Lehner and Döll 2004; Olefeldt et al. 2021) to upscale field plot-based emissions using generalized landcover types linked to *in situ* flux measurements (Matthews and Fung 1987; Kuhn et al. 2021; Kou et al. 2022), which allows for error estimates derived from statistical models and distributions (Davidson et al. 2017; Kou et al. 2022; Ramage et al. 2024). However, landcover-class scaling often misses temporal responses related to intra or inter annual responses. Biogeochemical models that incorporate abiotic (climate, soil inundation dynamics) and biotic (plant, microbial) drivers of emissions have the ability to capture temporal variability to improve annual emission estimates and predictions (Zhang et al. 2023), however these models also require generalized landcover categories to simplify the landscape (Treat et al. 2018). Landcover-class scaling has rarely been coupled with paired field measurements of isotopes or microbes (Ganesan et al. 2018). Thus, a fundamental question remains: do generalized landcover types adequately represent the relationships between the above-(vegetation) and below-(microbial) ground processes that control climate-warming gas emissions?

The Arctic is a consequential and difficult region in which to address emissions scaling. It is projected to be a source of significant climate feedbacks (Schuur et al. 2015), and is poorly captured in most global earth system models (Canadell et al. 2025), yet rapidly changing conditions (Rantanen et al. 2022), high heterogeneity (Bartsch et al. 2023), and limited direct measurements (AMAP 2015), make it difficult to accurately estimate emissions. Methane is a key mediator of these feedbacks (Saunois et al. 2020; Schuur et al. 2022) and its emissions are increasing due to warming (Yuan et al. 2024) and permafrost thaw (Varner et al. 2022; Turetsky et al. 2020). Thaw drives transitions in plant and microbial communities (Standen and Baltzer 2021; Ernakovich et al. 2022), shifting methane production and consumption processes (Deng et al. 2017), fluxes (Kuhn et al. 2021), and isotopic signatures (McCalley et al. 2014). Understanding the current magnitude and response of Arctic methane emissions and related isotopic signatures is critical to constraining global carbon models (Saunois et al. 2020; Basu et al. 2022) and informing policy decisions (Natali et al. 2021). At an intensively-studied permafrost peatland ecosystem undergoing rapid thaw, we tested the consistency of above-and below-ground community types, the uncertainty of remotely sensing them, and the consequentiality of modeling their interconnection to upscaling climate feedbacks.

First, to test the consistency of above and below-ground communities, we characterized plants and microbiota across Stordalen Mire, Sweden (68°21′ N 18°49′ E, Supp Fig S.1), at sites representing the three main thaw stage habitats (Fig 1A; palsas are unthawed, bogs are partially thawed, and fens are fully thawed), including long-term measurement sites with more than two decades of autochamber observations of carbon gas fluxes (Holmes et al. 2022). We found a strong and significant correlation of belowground microbial communities with aboveground vegetation community composition. Each habitat’s microbiota was statistically distinct (p<0.001; Fig 1B; community divergence also weakly increased with geographic separation in some cases, see Supp Fig S2A, B). Any given sample’s microbiota was predictive of the thaw stage with 85-92% accuracy (random forest classifications identified twelve predictor lineages, including one methanogen, Supp Fig S3). The Mire-wide belowground microbial communities were strongly correlated with aboveground plant communities, significantly at all depths (Fig 1C; Procrustes correlations 0.75-0.84, p=0.001; Mantel correlations 0.54-0.66, p=0.001 Table S1; correlation was strongest at the surface, potentially reflecting the depth pattern of plant influence via roots and litter). This previously unreported relationship for Arctic ecosystems (and only minimally in other systems for any plant-microbial community beta diversity comparison across landscapes; e.g. Prober et al. 2015) established the potential for generalized landcover types to adequately represent belowground communities in permafrost peatlands. Our findings suggest that readily observable and remotely sensible vegetation patterns could serve as proxies for belowground microbial communities, potentially simplifying landscape-scale modeling of biogeochemical processes in rapidly changing Arctic environments.

**Figure 1:**
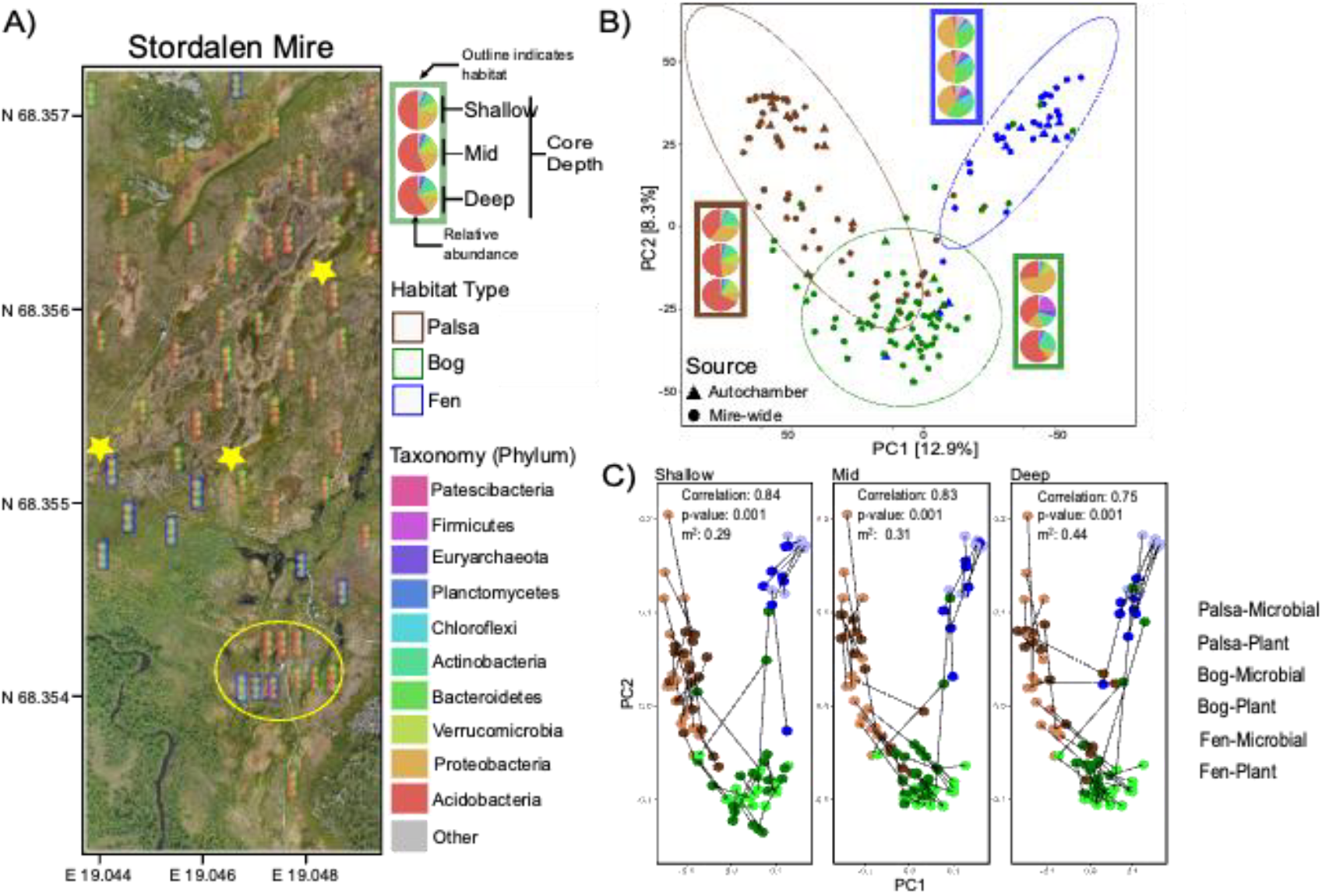
Thaw-stage habitats harbor distinct microbiomes, at the landscape scale, which co-vary with aboveground plant community composition. A. Microbiomes were sampled across a heterogeneous permafrost peatland landscape from the three thaw-stage habitats, at three depths (of 1 cm intervals, shallow average = 2 cm, mid = 11.6 cm, deep = 19.4 cm). Small overlain pie charts show top 10 microbial phyla (from 16S rRNA amplicon sequencing), and habitat type is denoted by the color of rectangular outline (categorized during sampling, and confirmed via aerial imagery, the latter shown as map). Sites denoted by yellow stars correspond to the example palsa, bog and fen pie charts shown in B. Yellow circled area surrounds an autochamber system and has been intensively studied over a decade; its observations have been used to validate biogeochemical models for this site’s habitat types, and are denoted by “Autochamber” in Figures 2B and 3B. B. Microbiome composition was significantly different (by PERMANOVA) among the thaw-stage habitats; ellipses indicate 95% confidence intervals, and inset pie charts are from the 3 starred sites in A. This pattern is similarly recapitulated by NMDS, Fig S4. C. Superimposed ordinations of aboveground plant (lighter colors) and belowground microbial (darker colors) community composition for peat from each sampled depth, with paired samples connected by lines, show a strong and significant correlation of above-and belowground communities, regardless of depth (via Procrustes analysis; “correlation” is a measure of similarity between community structures, m^2^ is a metric of the goodness-of-fit).

To test the relevance of this aboveground/belowground linkage for scaling methane emissions, we investigated how the methanogenic and methanotrophic communities, methane emissions, and methane isotopic signature varied by habitat (Fig 2, Fig S5). Consistent with the overall microbial community (Fig 1C), the structure of the methanogenic community was correlated with that of the aboveground plant community (p=0.001 at all depths, correlation ranged from 0.75 at 1-5 cm to 0.47 at 20-24 cm; Fig 2A), as was the methanotrophs’ (p**≤**0.002, correlations 0.42-0.56; Fig S5). Methanogens’ compositional divergence with geographic distance across the landscape was even weaker than in the community overall, and restricted to the fen (Fig. S2C versus S2B), potentially underscoring the importance of understanding hydrologic connectivity and dispersal as the fen habitat continues to expand, both at Stordalen and across the Arctic (Varner et al. 2022; Turetsky et al. 2020). In addition, the communities’ potential for the production of methane (as relative abundance of the taxonomically inferred methanogenesis pathway; see Methods) was statistically different between habitats (p<0.001) and increased across the thaw gradient (Fig 2B).

**Figure 2.**
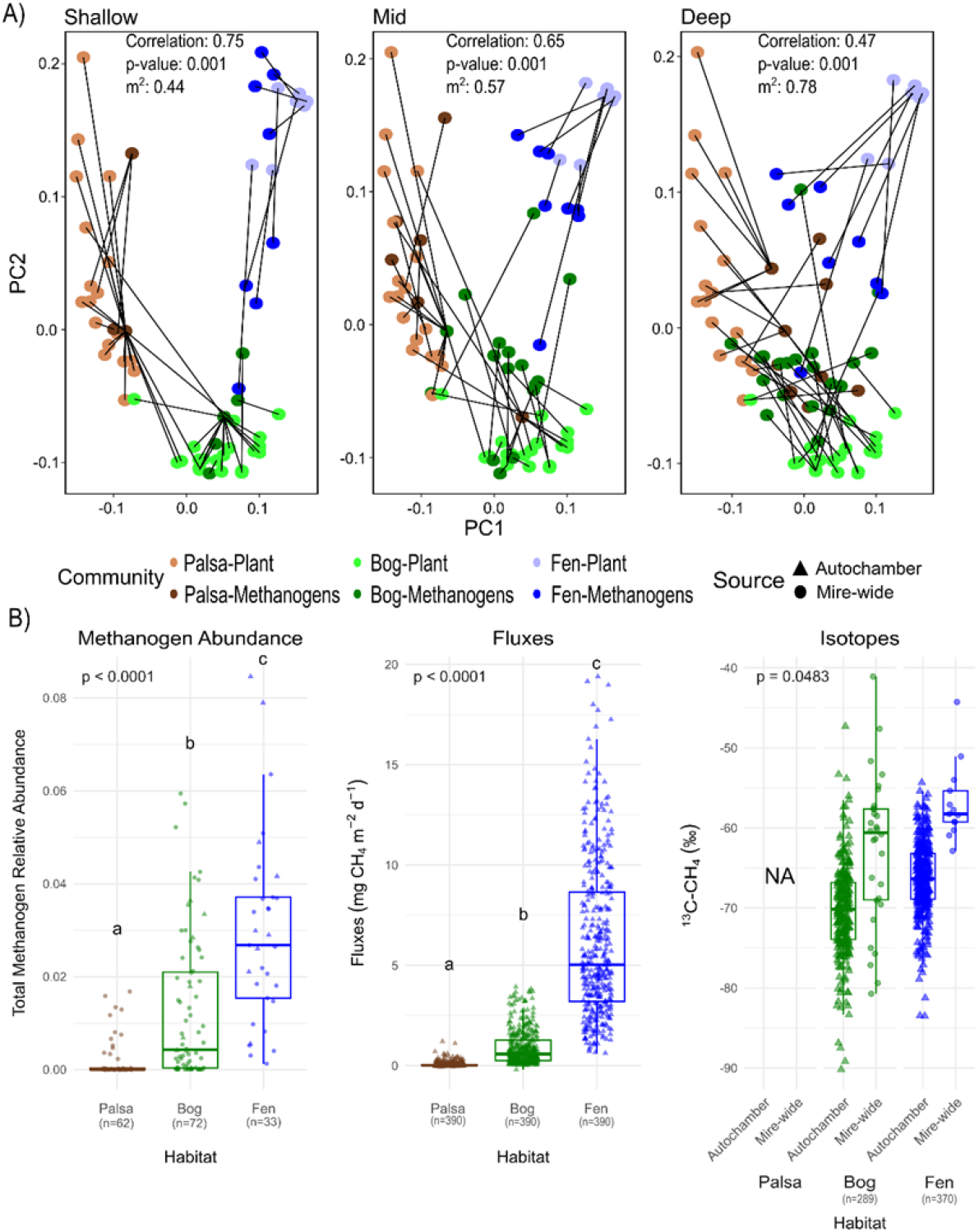
Aboveground/belowground consistency extends to methane cycling. A. Superimposed ordinations of community compositions of aboveground plants (lighter colors) and belowground inferred methanogens from each sampled depth (darker colors; via Procrustes, see Methods; left to right panels shallow to deep), with paired samples connected by lines, show strong and significant correlation (decreasing with depth) of inferred methanogen and plant communities. B. The inferred relative abundance of the methanogenesis pathway (as a portion of total inferred metabolic pathways, see Methods) in the microbiome differed significantly among the three habitat types, increasing with thaw stage (first panel), along with the flux of emitted methane (second panel, average daily fluxes per habitat from autochambers May-Nov 2015, averaged for each habitat), which becomes isotopically heavier (from autochamber fluxes and mire-wide porewater samples, third panel). Significant differences among habitats were assessed by Kruskal-Wallis test (methanogenesis), one-way ANOVA for log transformed fluxes, and t-test for emitted and porewater isotopes; for isotopes, these are shown separately for porewater and fluxes due to the known offset between them (e.g. Kuhn et al. 2024). Pairwise differences between habitats are indicated by letters above (a, b, c) and were assessed by a Dunn test with Benjamini-Hochberg (BH) multiple comparison correction for methanogenesis and Tukey Post-Hoc Test for log-transformed fluxes. Boxes indicate the interquartile range (IQR), with the median denoted by horizontal internal lines, and whiskers indicate 1.5 times the IQR.

To test for differences in methane emissions among thaw stages, we then leveraged the high temporal-resolution autochamber measurements since the spatially-resolved landscape-scale sampling was from a single time point, and captured methane porewater concentration and ^13^C isotope measurements (not fluxes), and due to fluxes’ high temporal variability (McCalley et al. 2014), we then leveraged the high temporal resolution autochamber flux and isotope observations to test for differences among thaw stages. Supporting these integrated analyses of the landscape-scale and autochamber observations for connecting microbiota to emissions, the autochamber site microbial communities were within the distribution of those at the landscape scale, for both the overall communities and the inferred methanogenic pathways (Fig. 1B, 2B; the within-habitat variance was also not statistically different for bog or fen, Fig. S7). The thaw stage-associated increase in the microbial production potential of methane was reflected in a corresponding increase in autochamber methane fluxes (Fig. 2B). The ^13^C of methane fluxes increased across the bog-to-fen transition (Fig. 2B; fluxes are minimal from palsas), consistent with porewater isotopes across the landscape (Fig 2B). This pattern reflects a shift from hydrogenotrophic to increasingly acetoclastic production from bog to fen (McCalley et al. 2014), underlain by an increasing abundance of acetoclastic methanogens which are also significantly correlated to habitat at the landscape scale (p<0.02; Supp Fig. S6).

Further supporting landcover type-based inferences of methane dynamics, porewater isotopic signature was significantly correlated to total sedge leaf area (p=0.004; Table S2), which is a metric used in landcover classification schemes and can be derived by higher resolution remotely sensed image data (Olefeldt et al. 2021; Kuhn et al. 2024). Collectively, these findings demonstrate significant consistencies among: aboveground plant community composition, overall belowground microbial composition at all measured peat depths, relative abundance of inferred methanogenesis, and methane emissions and isotopes. This robust relationship supports the use of vegetation-based landcover classifications for inferring belowground processes and methane dynamics in permafrost peatlands.

Having established these aboveground-belowground connections, we then used remote sensing of vegetation to scale methane emissions to the landscape. Machine learning was used on WorldView-2 satellite images of the Mire (Fig 3A; 69.7 ha) to classify pixel profiles into vegetation cover classes (as in Palace et al. 2018) (Fig 3B), yielding a misclassification of 3.5%. We then applied a landcover-class scaling approach, assigning averaged autochamber flux observations for each habitat to the pixel-by-pixel habitat classes (Fig 3C). However, since emissions are highly spatiotemporally variable (McCalley et al. 2014; Varner et al. 2022), using bulk landcover-class averages likely misestimates landscape emissions. We therefore applied a mechanistic approach to scaling emissions via the Wetland-DNDC biogeochemical model (Deng et al. 2017), which incorporates spatiotemporal variation in weather, soil variables, and water table as well as above-and below-ground linkages. The model was previously validated for the Stordalen Mire autochamber site with high accuracy for methane fluxes and isotopes. During the validation, the discrepancies between the simulations and observations were less than 20% for the seasonal total methane fluxes and 1‰ for the ^13^C isotopic signature in methane for both the bog and fen sites (Deng et al. 2017). A prerequisite to its application at the landscape scale was generating spatiotemporally dynamic water table estimates (Fig 3D) as input. This was accomplished by fusing processed time series observations of the Phased Array–type L-band Synthetic Aperture Radar (PALSAR) from the Advanced Land Observing Satellite (ALOS-2), with *in situ* sensors that captured hourly measurements of water height (Persson et al. 2012) (see Methods). The model then provided a dynamic portrait of methane fluxes and isotopes across the Mire (Fig 3E and F).

**Figure 3.**
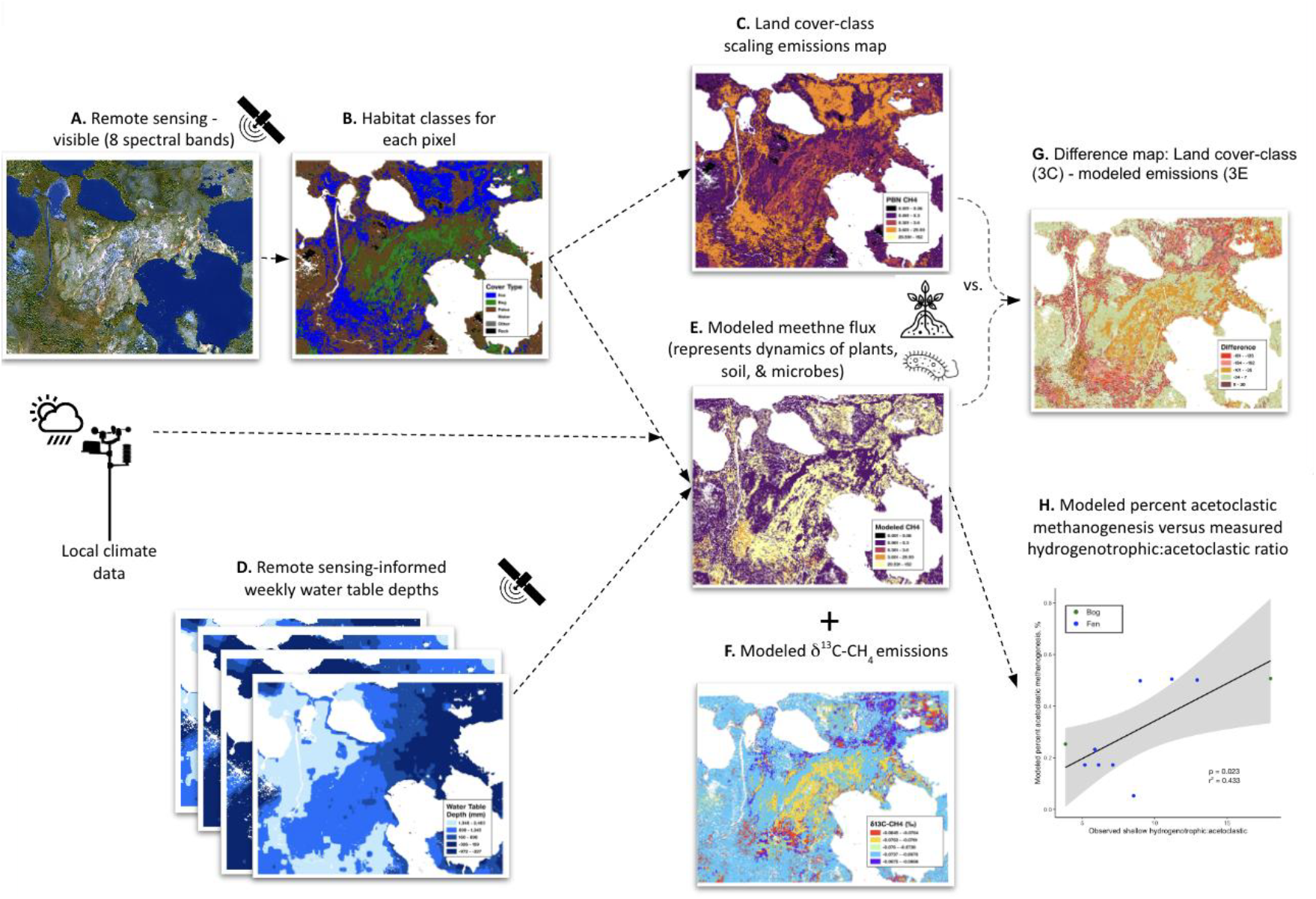
Remotely-sensed above-ground plant communities and water table can be used to estimate methane and isotopic (^13^C-CH_4_) emissions at the landscape scale. A. Remote sensing image of Stordalen Mire, Sweden (WorldView-2 satellite). B. Supervised classification of cover type, including 3 thaw-stage habitats. C. Methane emission across the landscape calculated via a landcover-class scaling approach using 2014 autochamber data. D. Water table depth using remote sensing (PALSAR sensor, ALOS-2) calibrated by field measurements. E. Methane emission across the landscape calculated via biogeochemical modeling (Wetland-DNDC), driven by B and D and climate data. F. Methane isotopic composition (δ^13^C-CH_4_ ) of emissions from the model in (E) G. The difference between (C) and (E); inset: the range of difference values for each habitat class. H. The modeled percent acetoclastic methanogenesis vs. observed hydrogenotrophic:acetoclastic ratio in the shallow peat samples (for samples with observed acetoclastic methanogen lineages).

Modeling resulted in markedly higher Mire-wide emissions than the static landcover-class scaling approach (∼2.5X, 56.2 vs. 19.8 mg methane m^-2^ d^-1^, Figure 3G), with the largest discrepancies in fens and the smallest in palsas (i.e., where it is most consequential to methane emissions estimates; inset Figure 3G). The isotopic signatures also differed; modeled Mire-wide flux-weighted annual isotope emissions are ^13^C depleted compared to landcover-class scaling approach (-73.0 ± 3.0 ‰ vs. -68.6 ± 1.0 ‰). Thus, integrating the additional complexity of dynamic water table, climate, and aboveground-belowground linkages produced a higher and isotopically lighter (implying less cumulative acetoclastic production and/or oxidation) flux estimate for the region. Here, all model outputs of belowground methane processes (gross production and its pathways, oxidation, net flux and ^13^C signature) were significantly (p<0.05) correlated to observed plant, geochemical, and microbial field metrics (Tables S2 and S3, Fig 3H). This stands in contrast to simple landcover-class scaling which cannot be correlated point by point to heterogeneous field observations as it applies a single value to each habitat. Methane emissions can therefore be consistently and accurately simulated by incorporating remotely sensed plant community proxies of microbial methane-cycling community structure into seasonally and system-integrated biogeochemical models in these rapidly thawing systems.

Atmospheric methane is rapidly increasing and becoming isotopically lighter (^13^C-depleted; Nisbet et al. 2023) and while the underlying causes are unresolved (Turner et al. 2019), they appear to be biological (Oh et al. 2022). Resolving methane source contributions requires scaling local ground-based measurements to regional fluxes, with high latitudes a priority due to their increasing biogenic methane emissions (Varner et al. 2022; Yuan et al. 2024). In addition, the distinct emissions profiles of high-latitude bogs and fens (e.g. at Stordalen Mire, a ∼6 ‰ difference in Fig 2B in 2015, and a ∼14 ‰ in 2011 from McCalley et al. 2014, and a ∼12 ‰ difference across five circumpolar peatlands in Kuhn et al. 2024) warrant their distinct representation in scaling. Here, habitat-explicit upscaling of emissions’ isotopic signatures at Stordalen Mire’s produced landscape values appreciably different from the single “wetland” habitat values used in top-down atmospheric model inversions (here ∼-73 ‰ via modeling and -69 ‰ via landscape cover scaling, versus ∼-60 ‰; Bousquet et al. 2006), a difference reflective of permafrost-associated wetlands generally (Fisher et al. 2017; Oh et al. 2022). This inaccuracy is increasingly consequential to accurate source partitioning (McCalley et al. 2014; Kuhn et al. 2024) as high-latitude systems diverge faster and further from their baseline emission states (IPCC, 2021). Collectively this work demonstrates that habitat-explicit scaling of belowground methane processes across heterogeneous landscapes, via remote sensing and modeling, is not only feasible but produces more accurate methane source profiles. Given methane’s potency as a climate-active gas, accurate scaling from the archaea that produce it to the atmosphere that holds it for 9.25 years (Short-lived Climate Forcers 2023; IPCC, 2021) is consequential to understanding and predicting ongoing biosphere-climate feedbacks.

How consistently are strong aboveground-belowground methane scaling relationships observed across ecosystems and timescales? Further research should test other climate-consequential habitats such as seasonally-inundated tropical forests (Melack and Hess 2023). In addition, since aboveground-belowground connectivity timescales differ among ecosystems and by rate of environmental change, how does scaling timescale impact scaling validity? In Stordalen Mire, where habitat conversion is occurring rapidly (Varner et al. 2022), this single-year study and one spanning 7 years (Cronin et al. 2025) suggest that a bog is a bog and a fen is a fen for methane emissions scaling purposes.

## Collective Author Designations

### The Archaea to Atmosphere (A2A) Project Team

Eleanor E. Campbell^1^, Justin Fisk^2^, Cheristy Jones ^1,3^, Jamie Lamit ^4^, Apryl Perry ^1,3^, Clarice R. Perryman ^5^, Joanne Shorter ^6^, Franklin Sullivan^1^, Peter Tansey ^1,7^, Nathan Torbick^1,8^, Beth Ziniti^9^

1. Earth Systems Research Center, Institute for the Study of Earth, Oceans and Space, University of New Hampshire; Durham, USA.
2. TBD
3. Department of Earth Sciences, University of New Hampshire; Durham, USA.
4. Department of Biology, Syracuse University, Syracuse, USA
5. Department of Earth System Science, Stanford University, Stanford, USA
6. Aerodyne Research, Inc., Billerica, USA
7. Department of Natural Resources and the Environment, University of New Hampshire, Durham, USA
8. Mitti Labs, Lee, USA
9. Regrow Ag, Durham, USA

### EMERGE Institute Coordinators

Jessica Ernakovich^1^, Maria Florencia Fahnestock^2,3^, Regis Ferriere^4^, Michael Ibba^5^, Matthew B. Sullivan^6,7,8^, Malak Tfaily^4^, Ben J. Woodcroft^9^, and Ahmed Zayed^6^

1. Department of Natural Resources and the Environment, University of New Hampshire, Durham, USA
2. Department of Earth Sciences, University of New Hampshire; Durham, USA.
3. Earth Systems Research Center, Institute for the Study of Earth, Oceans and Space, University of New Hampshire; Durham, USA.
4. Department of Ecology & Evolutionary Biology, University of Arizona; Tucson, USA.
5. Schmid College of Science and Technology, Chapman University, Orange, USA
6. Microbiology Department, The Ohio State University; Columbus, USA.
7. Center of Microbiome Science, The Ohio State University, Columbus, USA.
8. Department of Civil, Environmental, and Geodetic Engineering, The Ohio State University, Columbus, OH, USA
9. Centre for Microbiome Research, School of Biomedical Sciences, Queensland University of Technology (QUT), Translational Research Institute, Woolloongabba, Australia

## Data and materials availability

All data used in this study (with exceptions for proprietary datasets) are available and linked from https://emerge-db.asc.ohio-state.edu.

